# Single-cell analysis of an adult IBD INCEPTION cohort reveals Galectin-linked disease mechanisms

**DOI:** 10.64898/2026.06.30.735473

**Authors:** Matthew Leipner, Peter Rimmer, Samantha Tull, Alexandra Paun, Virginie Sandrin, Jenefa Begum, Adel Abo Mansour, Anella Saviano, Naveen Sharma, Jonathan Cheesbrough, Francesco Maione, Patrick Trenkle, Achim Klein, Sabrina Danilin, Tariq H Iqbal, Asif J Iqbal, Daniel Regan-Komito

**Affiliations:** Computational Sciences Center of Excellence, F. Hoffmann-La Roche, Ltd. Basel, Switzerland; Department of Gastroenterology, University Hospitals Birmingham NHS Foundation Trust, Birmingham, United Kingdom; Department of Microbes, Infection and Microbiomes, College of Medicine and Health, University of Birmingham, Birmingham, United Kingdom; Department of Cardiovascular Sciences, College of Medicine and Health, University of Birmingham, Birmingham, United Kingdom; Roche Pharma Research and Early Development, Roche Innovation Center Basel, F. Hoffmann-La Roche, Ltd., Basel, Switzerland; Department of Clinical Laboratory Sciences, College of Applied Medical Sciences, King Khalid University, Abha 62521, Saudi Arabia; ImmunoPharmaLab, Department of Pharmacy, School of Medicine and Surgery, University of Naples Federico II, Via Domenico Montesano 49, Naples 80131, Italy

**Keywords:** Inflammatory Bowel Disease, Crohn’s Disease, Ulcerative Colitis, Single-cell RNA sequencing, Galectin-9, Inflammatory monocytes

## Abstract

**Background and Aims:** The molecular pathogenesis of Inflammatory Bowel Disease (IBD) remains unclear. We aimed to establish a high-resolution immune landscape of treatment-naïve IBD to identify central drivers of disease onset and early pathogenic signalling.

**Methods:** We generated a single-cell atlas using intestinal biopsies from a large adult inception cohort of 137 individuals, including treatment-naïve Crohn’s disease (CD), ulcerative colitis (UC), and symptomatic non-IBD controls. We integrated scRNA-seq (1 million cells) with co-varying neighbourhood analysis (CNA) and unbiased tensor decomposition of cell-cell communication (CCC) networks. Findings were validated in vitro macrophage stimulation model and using serum from patients.

**Results:** The inception cohort exhibited significantly more homogenous compartmental diversity compared to benchmark reference studies (p < 0.001). Inflammation in both CD and UC was characterized by a marked expansion of inflammatory monocytes. Unbiased CCC analysis identified a dominant disease-specific signalling module centred on the Galectin family (LGALS1 and LGALS9). Galectin-9 expression was specifically enriched in inflammatory monocytes, which exhibited distinct transcriptional programs linked to antigen presentation and microbial sensing. In vitro, Galectin-9 acted as a potent stimulus, driving macrophages toward a pro-inflammatory phenotype. Clinically, serum Galectin-9 levels were significantly elevated in IBD patients and correlated with systemic inflammatory markers and treatment response.

**Conclusions:** Our data identify a galectin-monocyte signalling axis as a unifying inflammatory hallmark of early IBD. Galectin-9 serves as both a functional driver of mucosal inflammation and a dynamic biomarker, offering new opportunities for therapeutic targeting and disease monitoring from diagnosis.

## Introduction

Inflammatory bowel disease (IBD) comprising ulcerative colitis (UC) and Crohn’s disease (CD) results from an exaggerated immune response to an environmental trigger in genetically predisposed individuals (1). Recent epidemiological data confirms that the incidence of IBD is rising inexorably with the spread of industrialisation(2) and is particularly problematic for industrialised nations in the Northern hemisphere. For example, IBD is likely to affect up to 1% of the UK population by 2030 (2, 3). Genome wide association studies have implicated a broad range of risk genes relating to intestinal barrier function, host-microbiome interaction and the adaptive immune response (4, 5). Despite the advent of many treatments directed at various facets of the immune response there is currently no cure and even those interventions which initially work have an efficacy ceiling of ∼40% (6). This is likely due to tissue pathophysiological variation at baseline, variation in patient trajectories in response to different treatments, microbiome dysbiosis(7–10) as well as decreased drug exposure over time due to the development of anti-drug antibodies in some cases (11).

In recent years a number of studies have leveraged single cell transcriptomics to more deeply phenotype the intestinal mucosa of IBD patients(12–18). These studies have pointed to a number of different pathogenic signals including fibroblasts, myeloid cells and plasma cells for example. However, the majority of this work with a few exceptions (15, 17, 19) has involved patients with established disease who have been on various treatments or paediatric cohorts where monogenic mutations or single disease drivers are more prevalent. Given the known effects of various anti-inflammatory treatments not just on the intra-luminal milieu but also on the tissue architecture, identifying a central driver of disease initiation in these cohorts remains challenging.

In an effort to focus on the initiation of disease, intestinal biopsy samples were collected from a large cohort of newly diagnosed treatment naïve adult IBD patients attending the Birmingham IBD inception clinic (20). Colonic biopsies were taken at the time of the diagnostic colonoscopy. Intestinal biopsies from ‘symptomatic controls’ who were not deemed to have inflammation at endoscopy were also included as controls. The data establishes a robust interactome of the initiating inflammatory response in IBD onset and identifies several potential initiating signals including carbohydrate binding lectins among others.

## Methods

### 2.1 Patient Enrolment

Adult (≥18 years) patients with suspected IBD, based broadly on symptoms and elevated faecal calprotectin levels (no set threshold), were triaged to a rapid access IBD ‘inception’ clinic. Patients who agreed to be contacted regarding research were approached prior to investigation and recruited to the study (Bloomsbury London Research Ethics Committee, reference 21/PR/0515, IRAS number 287279) at the time of the first ‘new patient’ appointment. Patients who had previously received treatment for IBD, or other immune mediated inflammatory diseases, were excluded.

Serum samples were obtained at the first new patient appointment. Mucosal punch biopsies were obtained using biopsy forceps (as per standard of care) at the time of the index diagnostic colonoscopy from patients with CD, UC and symptomatic control. Region of biopsy was at the discretion of the endoscoping physician as dictated by the disease distribution. Where possible, region of sampling was matched between inflamed and non-inflamed sites, with 10 biopsies of each type sought. Inflammation was assessed by macroscopic appearance and confirmed by routine H&E histological analysis. Outside of cases where inflamed mucosa was confined to the rectum, this site was avoided for sampling due to anticipated differences in cell populations and gene expression (21, 22).Post diagnosis treatment was not protocolised and followed standard of care led by an IBD multidisciplinary team in a large tertiary referral centre. Clinical outcome data was collected prospectively with treatment responses determined against pre-determined criteria.

### 2.2 Healthy Blood donors

Human blood was collected from healthy donors in EDTA tubes (Griener bio-one) via venepuncture. Written informed consents were obtained from all donors as well as approval from the University of Birmingham Local Ethical Review Committee (ERN_12-0079C).

### 2.3. Isolation of peripheral blood mononuclear (PBMCs) cells

PBMCs were isolated from whole blood via density gradient centrifugation on a layer of Histopaque 1077 (Sigma). Briefly, 5ml of whole blood was layered on top of 5ml of Histopaque 1077 and centrifuged at 800g for 45 minutes at room temperature (RT) with no brake or acceleration. The PBMC buffy coat was collected and washed twice with ice cold MACS buffer (PBS without Ca^2+^ and Mg^2+^, 2mM ethylenediaminetetraacetic acid (EDTA; Sigma Aldrich) and 0.5% bovine serum albumin, (BSA; Gibco), before total cell counts were performed using the Cellometer Auto T4 (Nexcelcom Bioscience).

### 2.4 Culture and stimulation of monocyte-derived macrophages

Isolated monocytes (detailed protocol in Supplementary methods) were resuspended in pre-warmed 0.15% BSA in RPMI 1640 (gibco) and seeded (3.5 x10^5^ cells per 500µl) in a 24 well tissue culture plate. Cells were left to adhere for 1 hr at 37°C and 5% CO_2_ to eliminate contaminating cells, which was followed by replacement with fresh macrophage differentiation media containing RPMI 1640, 50ng/ml MCSF (Peprotech) and 10% FBS (Sigma). Cells were cultured for 6 days with fresh media replacements every 2 days (23). On day 6, macrophages were stimulated either with 100ng/ml Lipopolysaccharide (LPS; Sigma) alone, 100nM Galectin-9 (Biotechne) alone or in combination for 16 hours, after which cells were rinsed with PBS and lysed in 350ul RLT lysis buffer (Qiagen). RNA was extracted using the RNAeasy RNA extraction kit (Qiagen) according to the manufacturer’s instructions.

### 2.5 Bulk RNA sequencing

#### 2.5.1 Preparation of sequencing libraries

Following RNA isolation, RIN (RNA integrity numbers) was measured using the High Sensitivity RNA ScreenTape® (Agilent) prior to performing sequencing. All samples attained RIN scores >7 and were processed by University of Birmingham Genomics service. Library prep was performed using the Lexogen QuantSeq 3‘ mRNA-Seq Library Prep Kit FWD (Illumina) followed by quality checks carried out using the Agilent TapeStation DNA 1000 tape (Aligent) and DNA HS Qubit™ (Fisher Scientific). The final RNA sequencing libraries were sequenced using the NovaSeq 6000 (Illumina).

#### 2.5.2 RNA sequencing analysis

Sequencing data files were generated in fastq format, and read quality was assessed using the FASTQC package in RStudio. Trimming was performed using BBDuk to remove adapter sequences, poly-A tails, and low-quality bases. Post-trimming, FASTQC analysis was conducted again to evaluate improvements in read quality. Sequenced reads were mapped to the human genome using the STAR aligner. Prior to alignment, the genome index was generated from the *Homo_sapiens.GRCh38.dna.Primary_assembly* reference genome and the corresponding annotation file *Homo_sapiens.GRCh38.101*, both obtained from the Ensembl database. Gene-level read counts across all samples were quantified using FeatureCounts.

#### 2.5.3 Analysis of differentially expressed genes

Differential gene expression (DEG) and pathway analysis was all performed using the iDEP integrated web application for bulk RNAseq analysis. The gene counts table generated with featureCounts was uploaded to iDEP and subsequent normalisation and analysis was performed. Data was normalised using variable stabilising transformation (VST) and the default filter of 0.5 counts per million (CPM) was used to filter out genes with low read counts. DEG analysis was performed using the DESeq2 package integrated in the iDEP web application. Enrichment analysis of DEGs was performed based on Gene Ontology (GO) databases. All heatmaps, PCA and MA plots were generated with the iDEP online application. iDEP is freely accessible at: http://bioinformatics.sdstate.edu/idep/

### 2.6 Biopsy tissue digestion

Freshly obtained punch biopsies were immediately transferred to ice cold CryoStor® CS10 (Biolife Solutions, Bothwell, WA, USA). Samples were then transferred to a - 80°C freezer for controlled rate freezing in a CoolCell® (Corning, NY, USA). Samples were transferred to liquid nitrogen after 24 hours and maintained at -80°C through temperature-controlled shipping to Roche. Biopsies were rinsed in ice cold PBS and gently minced using a scalpel. Tissue digestion was performed in complete digestion buffer (ATCC-modified RPMI 1640 supplemented with 1 U/mL penicillin-streptomycin, 50 µg/mL gentamicin, 10 mM HEPES, and 2% FBS) containing 1 mg/mL Collagenase D (Sigma) and 100 µg/mL DNase I (Sigma). Samples were incubated at 37°C on a heated shaker (250 rpm) for two consecutive 30-minute intervals. The resulting cell suspension was quenched with FBS containing 30 mM EDTA, filtered through a 40 µm cell strainer, and washed. All subsequent steps were performed on ice. Cells were resuspended in ACK lysis buffer (Gibco) for red blood cell lysis, followed by an additional wash and resuspension in epithelial cell solution (HBSS supplemented with 1 U/mL penicillin-streptomycin, 10 mM HEPES, 10 mM EDTA, and 2% FBS) for 3 minutes. The cell suspension was then passed through a second 40 µm strainer to remove epithelial cells and resuspended in 0.04% PBSA for further processing.

### 2.7 Single Cell RNA Sequencing

#### 2.7.1 Preparation of sequencing libraries

Single-cell libraries were generated using the Chromium Next GEM Single Cell 3’ Reagent Kit v3.1 (10x Genomics). Libraries were fragmented, end-paired, A-tailed, and ligated with Illumina-compatible adapters, quality-controlled via Agilent Bioanalyzer, and quantified by qPCR. Final libraries were sequenced using a paired-end configuration on an Illumina NovaSeq 6000 to a target depth of ∼50,000 read pairs per cell.

#### 2.7.2 Data Processing and Quality Control

Raw reads were aligned to the GRCh38-2024-A reference (GENCODE v40) using Cell Ranger v9.0.1 (24). Ambient RNA correction was performed using CellBender (25). Subsequent filtering was completed with Scanpy, removing cells with <100 UMI or <150 genes (26, 27). Cluster-specific outlier detection based on Median Absolute Deviation (MAD) was applied to UMI counts, gene counts, and mitochondrial content. Doublets were identified and excluded using scDblFinder (28).

#### 2.7.3 Annotation and Integration

Automated cell-type annotation was performed using CellTypist (29) at the sample level, utilizing gastrointestinal-specific reference models (22, 30). Samples were then merged and batch corrected by compartment, split between epithelial, immune, and stromal. Data integration and batch correction were performed using scVI (single-cell Variational Inference), incorporating mitochondrial and ribosomal percentages as covariates (31). Cell type annotations were then refined at the most specific level by manual curation using known marker genes. Dimensionality reduction and clustering were conducted via UMAP and the Leiden algorithm on the scVI latent space (32, 33).

#### 2.7.4 Co-Varying Neighbourhood Analysis (CNA)

To identify cell populations associated with inflammation, CNA was employed. This method identifies local neighbourhoods whose abundance varies with sample-level phenotypes while controlling for sex and technical batch. Here, inflammation status was used as the varying phenotype of interest. Statistical significance was determined via permutation testing (FDR < 0.05). Neighbourhood correlation scores (ncorr) were used to categorize cell populations as expanded or depleted in disease states (34).

#### 2.7.5 Cell-Cell Communication and Tensor Decomposition

Cell-cell communication analysis was performed using LIANA+, aggregating results across multiple ligand-receptor (LR) databases, including CellPhoneDB and OmniPath (35–38). To identify consistent communication patterns across the cohort, non-negative tensor factorization was performed using cell2cell with Tensorly (39) (http://jmlr.org/papers/v20/18-277.html). LIANA+ scores were formatted into a 4D tensor of shape [sample, LR pair, sender cell type, receiver cell type]. The optimal rank for decomposition was determined by elbow analysis, and factor activities were compared across disease conditions using Mann-Whitney U tests with Benjamini-Hochberg correction, grouping by CD-Inflamed, CD-Non-Inflamed, UC-Inflamed, UC-Non-Inflamed, and Non-IBD.

#### 2.7.6 Pathway Enrichment and Network Visualization

Biological interpretation of tensor factors was performed via Gene Set Enrichment Analysis (GSEA) on LR factor loadings using KEGG pathway annotations (40) with decoupleR (41). Additionally, pathway activity footprints were estimated using PROGENy (42). Communication networks were visualized by computing joint loadings of sender and receiver cell types, with significance thresholds applied to identify dominant signalling axes.

### 2.8 Serum Enzyme-linked Immunosorbent Assays (ELISA)

Standard ELISA protocols were performed for Gal-1, Gal-3, Gal-9, TNF, IL-6 and IL 23 on serum obtained from CD, UC and SC at baseline and longitudinally (Quantikine ELISA, R&D systems, Biotechne). https://doi.org/10.2307/4615733

## Results

### Single–cell profiling of intestinal biopsies across CD, UC, and symptomatic non–IBD controls

We generated a single-cell inception cohort using intestinal biopsies obtained from 137 individuals, including CD, UC, and symptomatic non-IBD controls (**Table 1**). Biopsies were processed through a harmonised pipeline involving cryopreservation, enzymatic dissociation, dead cell removal, and single cell RNA sequencing, yielding approximately one million high quality cells across 192 samples (**Figure 1a-c**). This design enabled resolution of mucosal cellular states across disease, tissue, and inflammation status

**Table 1:**
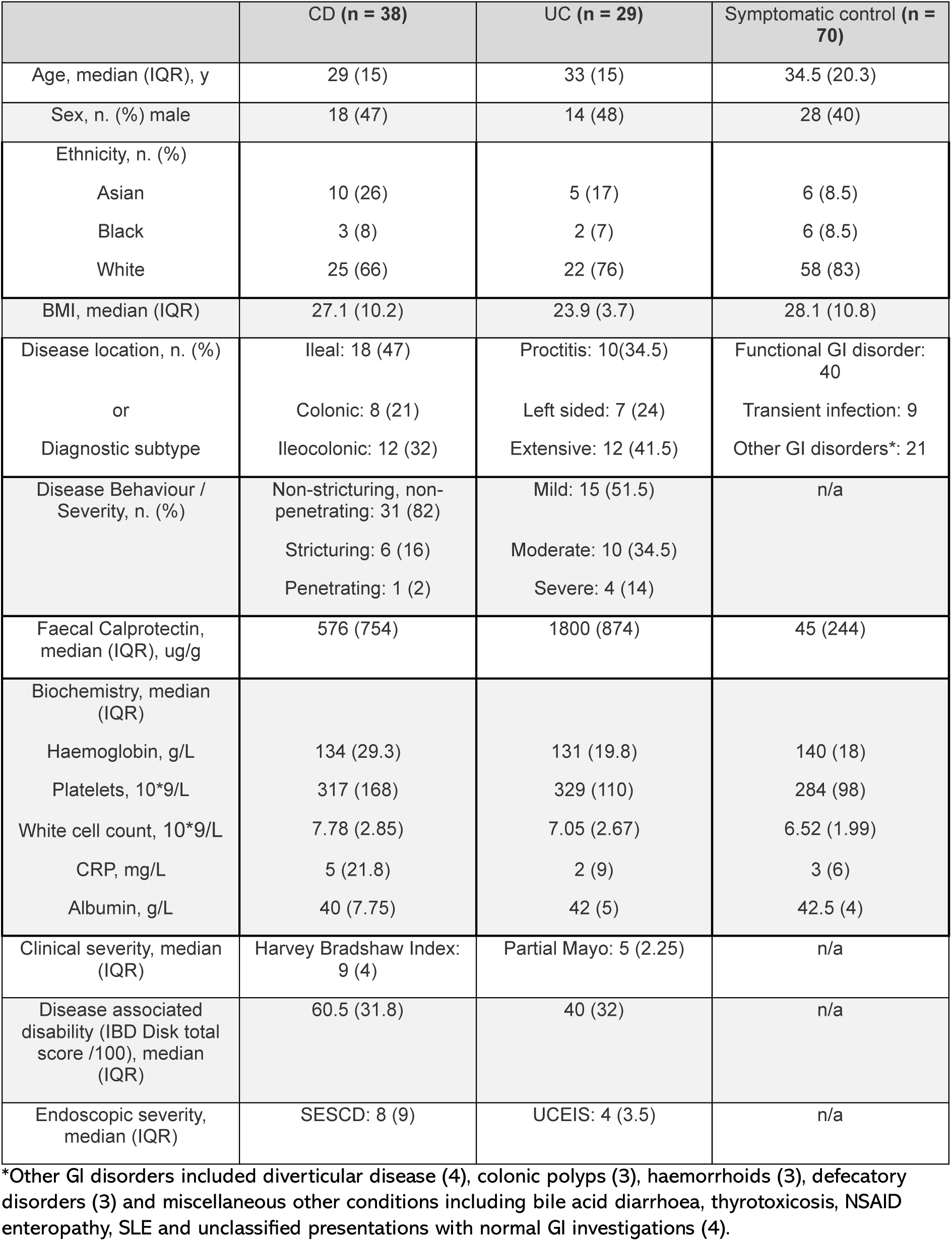
Patient demographics and baseline disease indices for treatment naïve IBD patients and symptomatic controls contributing samples to the scRNA seq dataset.

**Table 2:** Demographics and baseline disease indices for the patients contributing serum samples.

|  | CD (n = 69)<br>n. also in scRNA<br>seq dataset = 30 | UC (n = 59)<br>n. also in scRNA<br>seq dataset = 23 | Symptomatic<br>control (n = 20)<br>n. also in scRNA<br>seq dataset = 10 | Healthy control (n = 38) |
| --- | --- | --- | --- | --- |
| Age, median (IQR), y | 30 (17.3) | 35.5 (19.8) | 34 (15) | 26 (7) |
| Sex, n. (%) male | 25 (42) | 29 (58) | 10 (48) | 18(47) |
| Ethnicity, n. (%) |  |  |  |  |
| Asian | 18 (26) | 8 (14) | 1 (5) | 5 (13) |
| Black | 6 (9) | 5 (8) | 1 (5) | 1 (3) |
| White | 45 (65) | 46 (78) | 18 (90) | 32 (84) |
| BMI, median (IQR) | 24.5 (8.06) | 24.3 (4.6) | 29.6 (10.4) | 25 (5.2) |
| Smoking, n. (%) current<br>smoker | 11 (17) | 4 (7) | 3 (15) | Not known |
| Disease location, n. (%) | Ileal: 31 (45)<br>Colonic: 16 (23)<br>Ileocolonic: 22 (32) | Proctitis: 13(28)<br>Left sided: 12 (25)<br>Extensive: 22 (47) | n/a | n/a |
| Disease Behaviour /<br>Severity, n. (%) | Non-stricturing, non-penetrating: 53 (79)<br>Stricturing: 10 (15)<br>Penetrating: 4 (6) | Mild: 20 (43)<br>Moderate: 21 (45)<br>Severe: 6 (13) | n/a | n/a |
| Faecal Calprotectin,<br>median (IQR), ug/g |  |  |  | n/a |
| At referral | 676 (1210) | 1782 (1666) | 434 (447) |  |
| At time of serum<br>sampling | 516 (914) | 743 (1992) | 25 (131) |  |
| Biochemistry, median<br>(IQR) |  |  |  | n/a |
| Haemoglobin, g/L | 133 (25.5) | 135 (22.8) | 146 (19) |  |
| Platelets, 10*9/L | 345 (190) | 327 (145) | 281 (118) |  |
| White cell count,<br>10*9/L | 7.9 (3.51) | 6.95 (2.6) | 6.4 (1.8) |  |
| CRP, mg/L | 7.5 (13) | 2 (5.75) | 2.5 (4) |  |
| Clinical severity, median<br>(IQR) | Harvey Bradshaw<br>Index: 8 (5) | Partial Mayo: 4 (3) | n/a | n/a |
| Endoscopic severity,<br>median (IQR) | SESCD: 6 (8.5) | UCEIS: 4 (4) | n/a | n/a |

**Figure 1.**
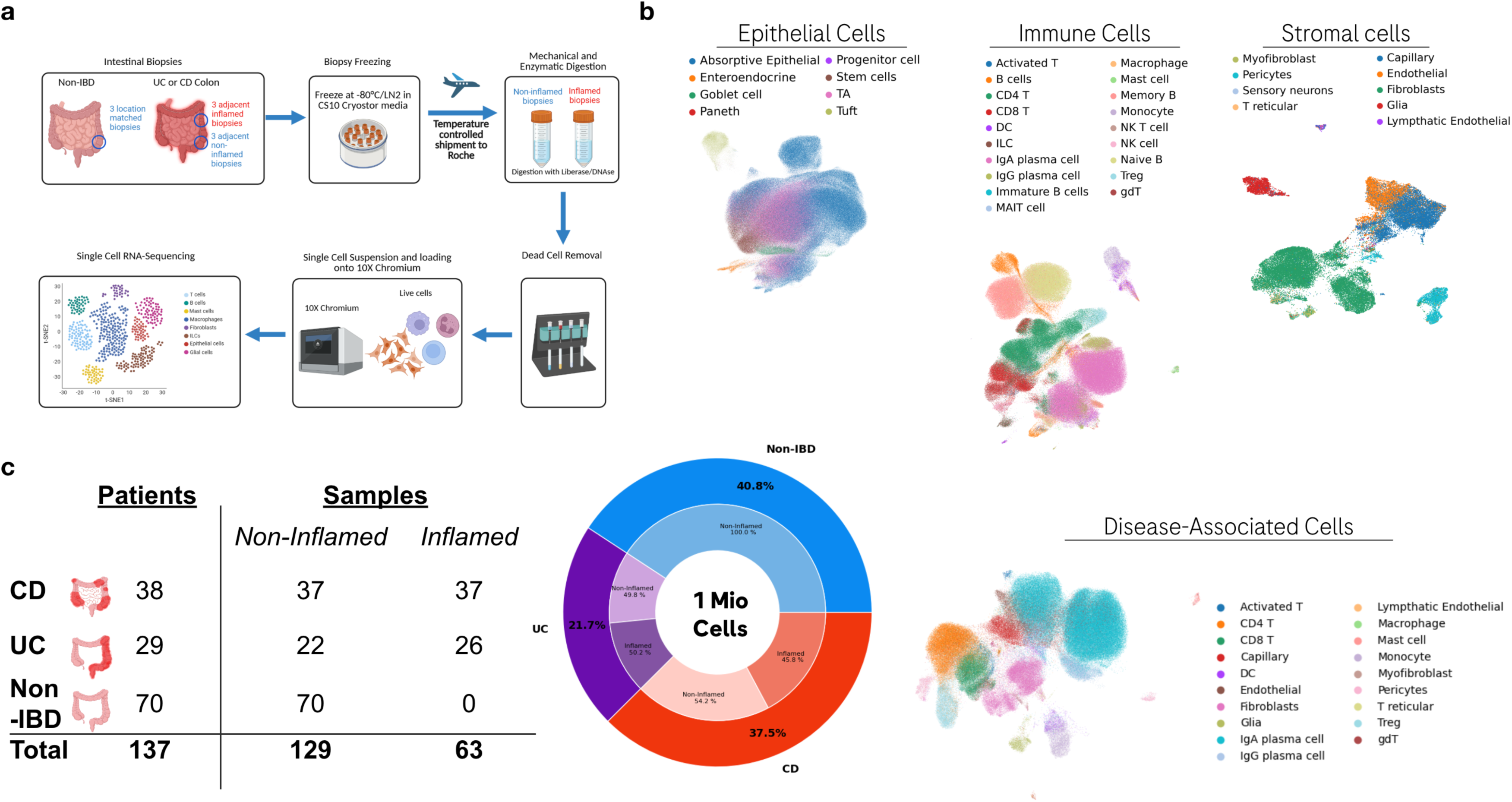
Single-Cell Inception Cohort of Colon and Ileum Biopsies from Healthy Individuals, Crohn’s Disease (CD), and Ulcerative Colitis (UC) Patients. A. Study design. B. Cell Type distribution, shown by Uniform Manifold Approximation and Projection (UMAP) embedding. C. Patient and sample counts within the dataset. D. Distribution of cells by disease and health status (inflamed vs. non-inflamed).

Unsupervised clustering identified 38 epithelial, immune, and stromal cell types, including absorptive and secretory epithelial subsets, diverse T and B cell populations, dendritic cells, monocytes, macrophages, fibroblasts, endothelial cells, and glial cells. The cohort included balanced representation across CD (37.5%), UC (21.7%), and non IBD controls (40.8%), supporting robust comparative analyses. Together, these data establish a high-resolution single-cell atlas of intestinal inflammation across the IBD spectrum in treatment-naive patients and symptomatic controls without inflammation.

### Disease associated cell states display altered abundance and reduced compartmental diversity in IBD

We next examined how inflammation alters mucosal cellular composition across CD, UC, and non-IBD samples. The dataset captures all major immune cell types associated with IBD, where marked differences are visible between disease and non-IBD controls. In CD, inflamed ileal and colonic tissues displayed marked expansion of monocytes, Th17 cells, Tregs, memory- and naive B cells, and class switching from IgA to IgG in plasma cells, with each population present at substantially higher proportions than in non-inflamed CD or symptomatic controls (**Figure 2a**). Among these, monocytes represented one of the most prominently enriched cell types, highlighting their central involvement in CD associated inflammation.

**Figure 2.**
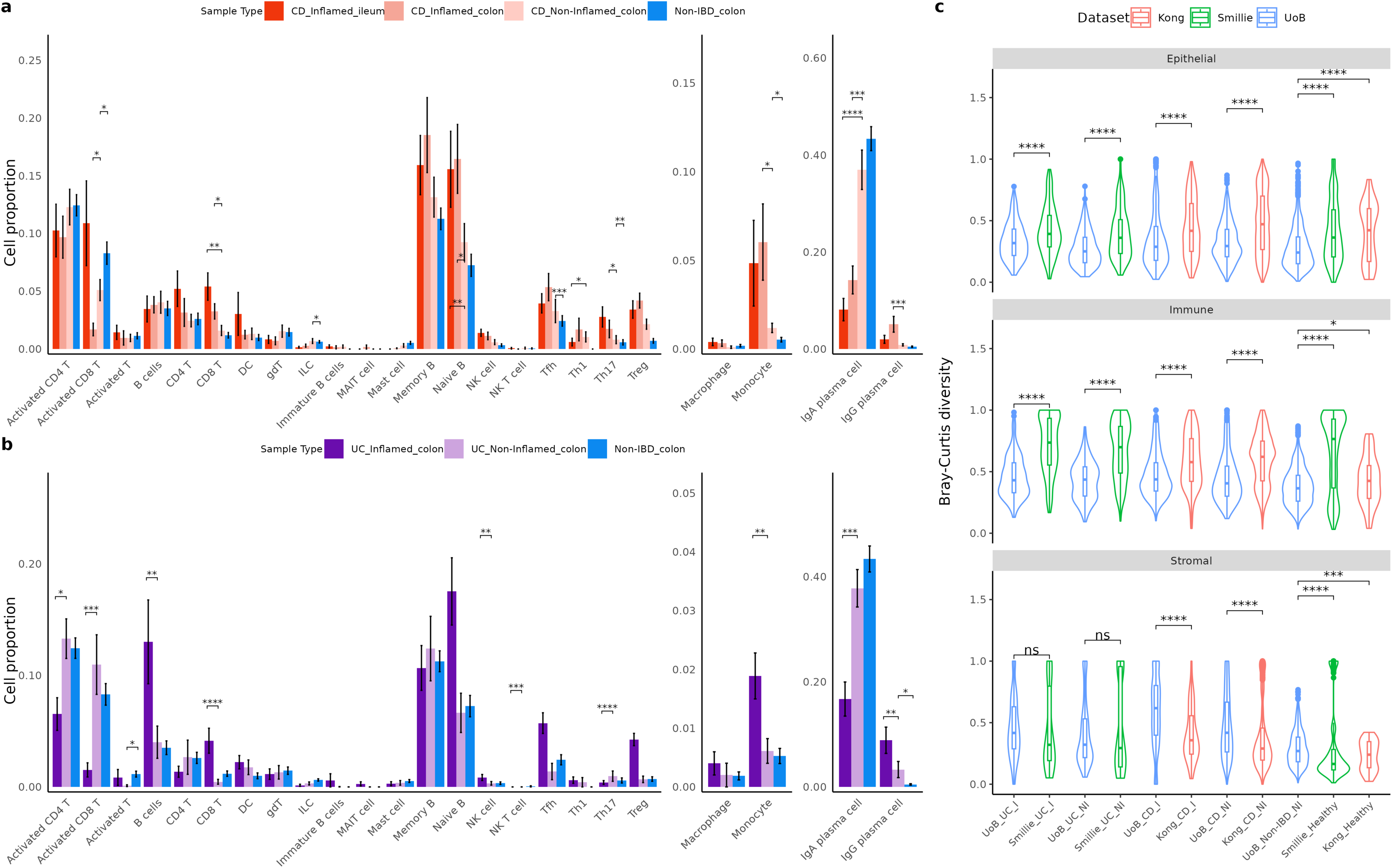
Distribution of cell types across disease and health status. A. Cell proportions of disease-associated cell types in Crohn’s Disease (CD). B. Cell proportions of disease-associated cell types in Ulcerative Colitis (UC). For each cell type, group differences were tested using two-sided Wilcoxon rank-sum tests. P values were adjusted by Benjamini–Hochberg (BH) within each cell type. C. Bray Curtis diversity within individual cell compartments comparing the Birmingham dataset with reference datasets. A lower Bray-Curtis diversity is representative of a more homogenous population. Bray–Curtis group comparisons were tested using two-sided Wilcoxon rank-sum tests for predefined dataset-matched contrasts; significance is displayed as star annotations.

UC inflamed samples displayed a distinct but overlapping immune signature, with elevated monocytes, Tregs, Tfh, mast cells, and IgG plasma cells compared with non-inflamed UC and non-IBD controls (**Figure 2b**). Notably, monocytes were again among the most proportionally expanded populations, underscoring their shared role across both major forms of IBD. These patterns position monocytes as a shared cellular hallmark of intestinal inflammation, while T and B cell driven responses diverge between CD and UC.

To assess intra-compartmental heterogeneity, we calculated Bray-Curtis diversity across epithelial, immune, and stromal compartments and compared the Birmingham dataset against published datasets (12, 16) (**Figure 2c**). Diversity scores were significantly lower across nearly all compartments, though particularly within the immune compartment (*P* < 0.001), indicating a more homogeneous representation of the treatment-naive, inflammatory cellular states. The Birmingham dataset showed strong concordance with reference cohorts, validating both the biological signal and technical robustness of the single cell framework. Moreover, this analysis highlights the increased robustness of our inception cohort compared to the increased biological and technical variability observed in previous studies.

### Unbiased cell-cell communication analysis reveals a Galectin centric signalling module activated during the onset of IBD

Disease-associated cell types were identified by co-varying neighbourhood analysis (CNA), and populations of inflammatory monocytes and fibroblasts were re-annotated based on known marker genes. To identify coordinated changes in mucosal signalling during inflammation, we first applied co-varying neighbourhood analysis to identify cell populations varying significantly with disease state. This analysis revealed strong enrichment of major epithelial, immune, and stromal populations in (CD) and (UC). Higher CNA loadings indicate greater relative presence of that cell type in inflamed IBD tissue compared with non-IBD. This analysis highlighted strong association of inflammatory monocytes, activated T cells, endothelial cells, and plasma cells in both CD and UC, establishing the cellular context in which communication networks are remodelled (**Figure 3a**).

**Figure 3.**
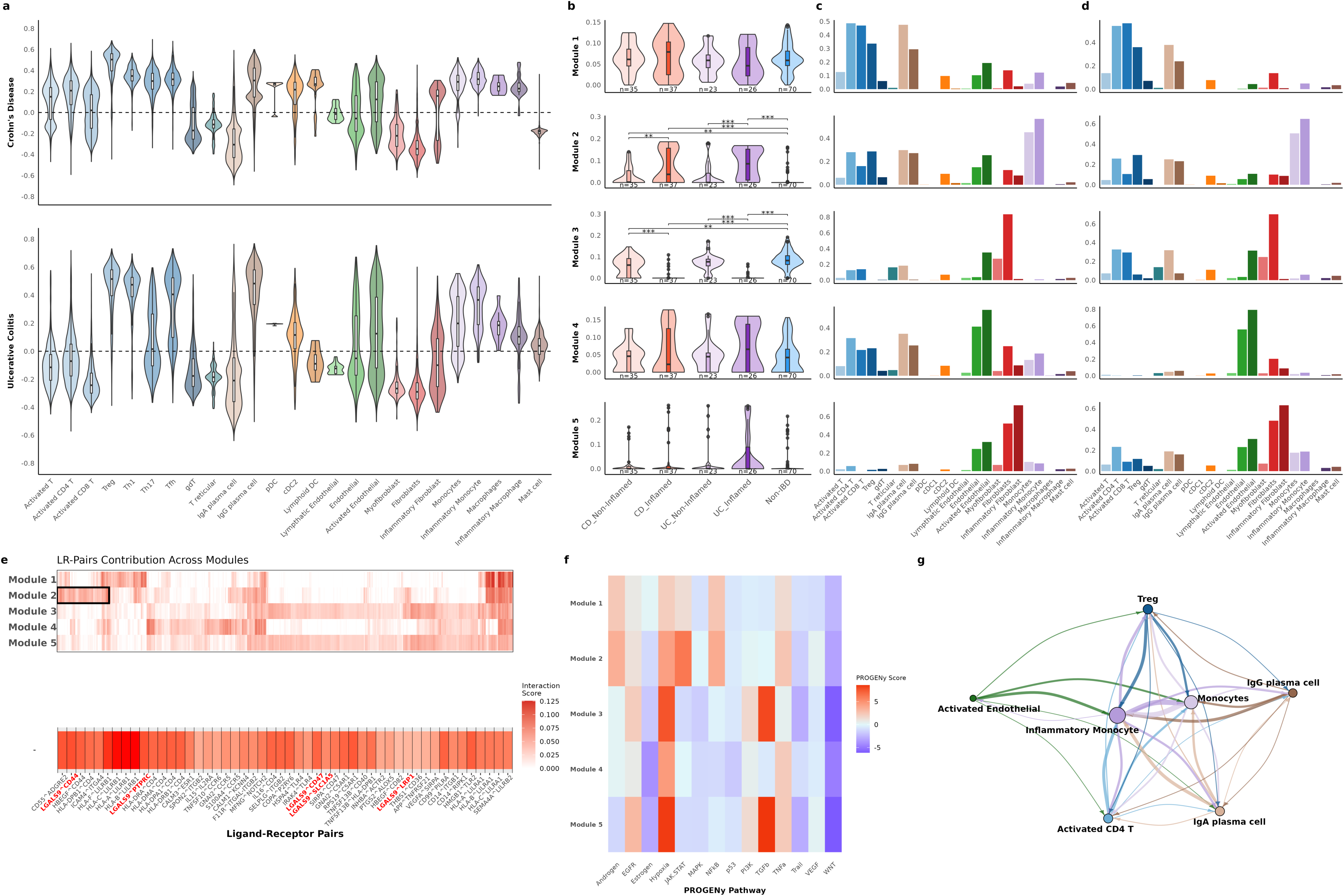
Cell-Cell Communication (CCC) within Disease-Associated Cell Types. A. Relative cell-type abundance inferred by co-varying neighbourhood analysis (CNA). Cell-type loadings indicate the relative enrichment of each population in inflamed disease samples compared with healthy reference samples. B. Overview of cell-cell communication (CCC) modules identified by tensor decomposition analysis. Heatmap depicts module weights across disease groups. Pairwise two-sided Wilcoxon rank-sum tests were performed between disease-state groups with BH correction. C. Sender-cell contributions to each CCC module. Higher weights indicate a greater contribution of a given cell population as a signalling source within the corresponding module. D. Receiver-cell contributions to each CCC module. Higher weights indicate a greater contribution of a given cell population as a signalling target within the corresponding module. E. Ligand-receptor interaction landscape across CCC modules. Heatmap shows ligand-receptor pair weights within each module, with the highlighted region indicating galectin-associated signalling interactions (LGALS1, LGALS3, and LGALS9) within Module 2. F. Pathway activity scores for each CCC module determined using PROGENy analysis. G. Network representation of active signalling interactions comprising Module 2. Nodes represent cell populations and edges represent predicted ligand-receptor-mediated communication pathways.

We next assessed cell-cell communication (CCC) using multiple well characterized methods and reference databases with the LIANA+ framework to unravel CCC signalling within each sample and further ran an unbiased tensor decomposition analysis on these CCC matrices using Cell2Cell to identify patterns associated with disease state. This analysis identified five distinct signalling modules, each representing a unique pattern of sender-receiver cell types and ligand-receptor (LR) interactions. Module scores (**Figure 3b**) demonstrated selective enrichment of modules 2 and 3 in inflamed CD and UC, indicating disease specific rewiring of mucosal communication networks.

Sender and receiver loadings (**Figure 3c & d**) revealed that inflammatory monocytes, activated endothelial cells, and IgA/IgG plasma cells were dominant contributors to these disease associated modules. Looking more closely at the ligand-receptor composition of these modules (**Figure 3e**) showed that module 2 was strongly enriched for galectin signalling, including LGALS1 and LGALS9 interactions with reported cognate receptors such as CD44, HAVCR2, and LRP1. Pathway activity analysis (**Figure 3f**) confirmed that this module shows increased canonical inflammatory cascades, including NFκB, JAK-STAT, and hypoxia pathways.

The structure of CCC signalling within module 2 (**Figure 3g**), revealed a tightly connected communication circuit linking inflammatory monocytes, activated CD4⁺ T cells, endothelial cells, and plasma cells, positioning galectin mediated signalling as a central, unbiasedly identified axis of inflammatory crosstalk in IBD.

### Galectin-9 expression is enriched in inflammatory monocytes within IBD tissues

To further interrogate galectin associated signalling across disease associated cell types, we mapped gene expression onto a UMAP embedding of the inflamed mucosal compartment. Clustering by cell annotations revealed distinct populations including inflammatory monocytes, inflammatory fibroblasts, activated T cells, and plasma cells (**Figure 4a**). When segregated by disease and inflammation status, these populations were predominantly derived from inflamed CD and UC samples, reinforcing their disease relevance (**Figure 4b**).

**Figure 4.**
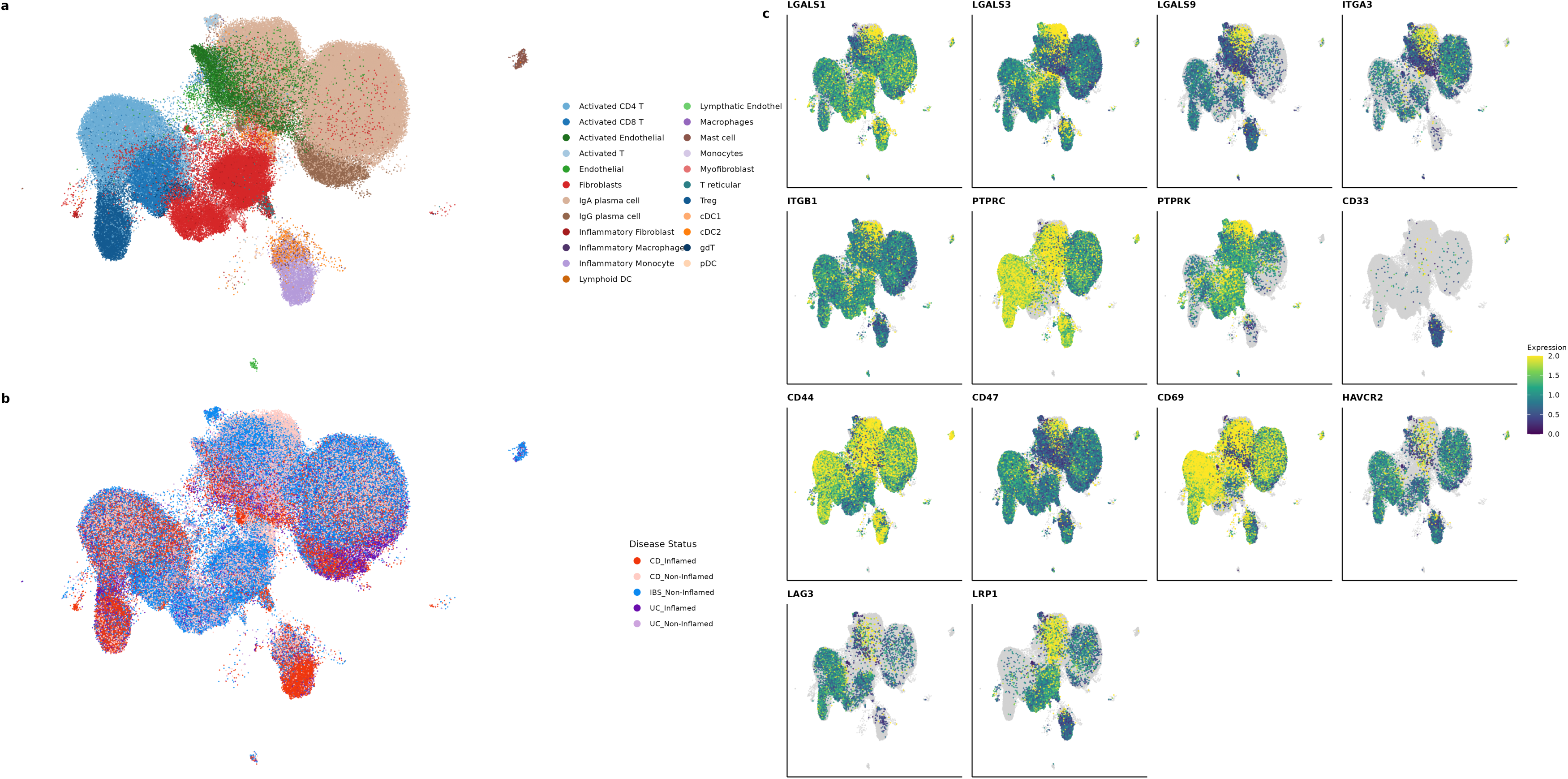
Galectin and Receptor Signalling within Disease-Associated Cell Types. A. UMAP of disease-associated cell types, coloured by cell type. B. UMAP of disease-associated cell types, coloured by disease and health combination.

Gene level overlays highlighted selective upregulation of galectin family members across specific cell types. Notably, LGALS9 was strongly expressed in inflammatory monocytes, suggesting a cell-intrinsic contribution to the galectin-centred communication module identified earlier. This enrichment was not observed in healthy or non-inflamed tissues, positioning inflammatory monocytes as key transmitters of galectin mediated signalling in the IBD microenvironment. Expression of galectin receptors and co signalling molecules (e.g., CD44, CD47, HAVCR2) further supported the presence of an active ligand-receptor axis within this niche (**Figure 4c**).

### Galectin positive inflammatory monocytes exhibit distinct transcriptional and pathway signatures

To dissect the functional heterogeneity within inflammatory monocytes, we stratified cells based on the expression of LGALS1 and LGALS9. Differential expression analysis revealed that LGALS1⁺ monocytes were enriched for genes involved in cytoskeletal organisation (e.g., S100A4, S100A6, ACTB) and inflammatory signalling (IL1β, FTH1), suggesting a pro-inflammatory and migratory phenotype (**Figure 5a**). KEGG pathway enrichment confirmed activation of antigen processing, oxidative phosphorylation, and ribosomal pathways, indicating heightened metabolic and immunological activity (**Figure 5b**).

**Figure 5.**
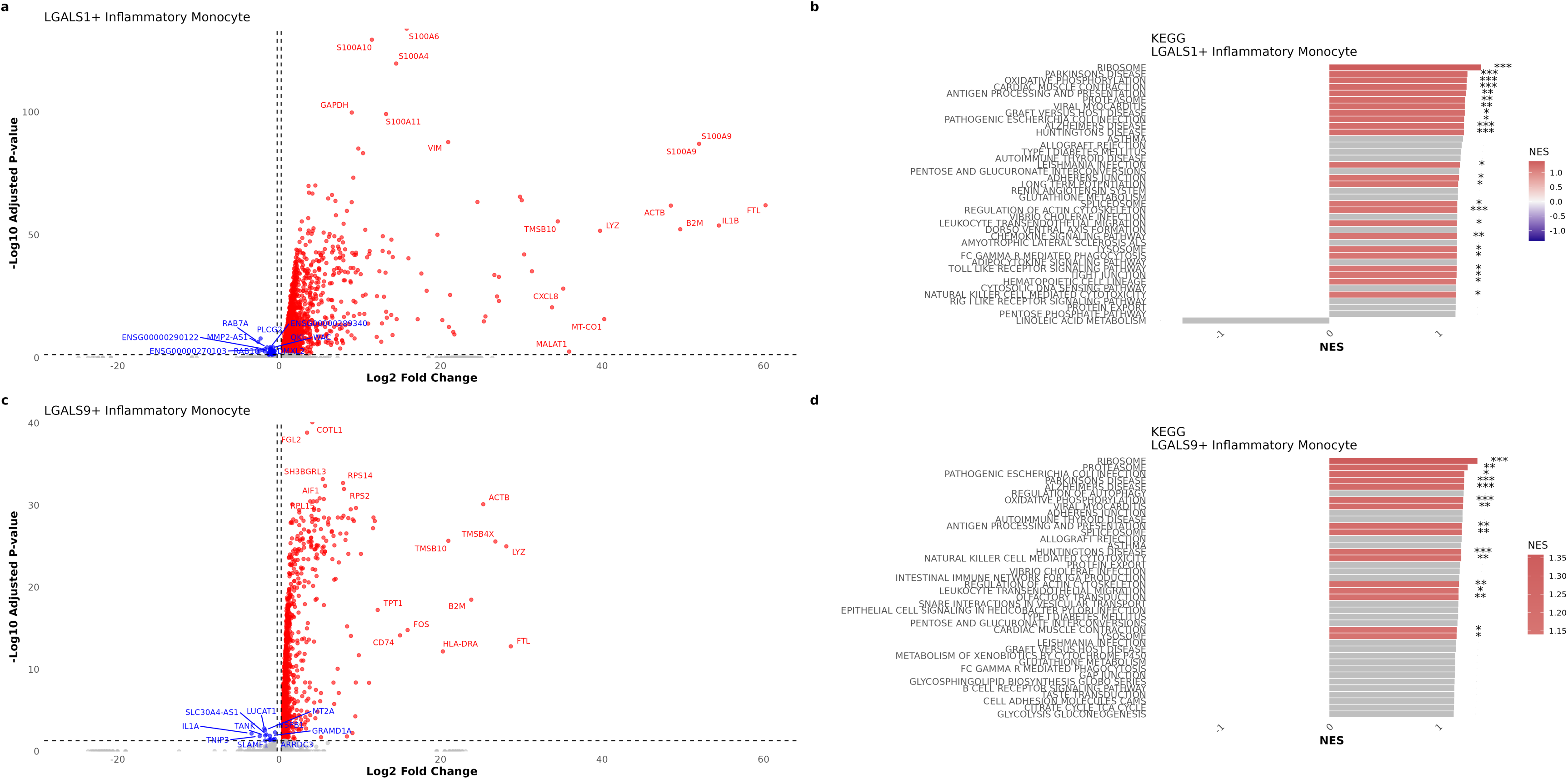
Profiling of Galectin-positive vs. Galectin-negative Inflammatory Monocytes. A. Volcano plot of Galectin-1 (LGALS1-) positive inflammatory monocytes vs LGALS1-negative inflammatory monocytes. B. KEGG enrichment of LGALS1-positive inflammatory monocytes, computed by FGSEA using ranked gene statistics; significance stars denote adjusted (p)-value thresholds. C. Volcano plot of Galectin-9 (LGALS9-) positive inflammatory monocytes vs LGALS9-negative inflammatory monocytes. D. KEGG enrichment of LGALS9-positive inflammatory monocytes, computed by FGSEA using ranked gene statistics; significance stars denote adjusted (p)-value thresholds.

LGALS9⁺ monocytes displayed a distinct transcriptional profile, with upregulation of FGL2, AIF1, CD74, HLADRA, and other genes linked to antigen presentation and immune regulation (**Figure 5c**). KEGG analysis highlighted enrichment of leukocyte migration, proteasome and ribosome pathways, alongside signatures associated with microbial sensing and neuroinflammatory processes (**Figure 5d**). These findings suggest that Galectin-1 and Galectin-9 mark transcriptionally and functionally divergent monocyte subsets, each contributing uniquely to the inflammatory landscape of IBD.

To assess the functional impact of Galectin-9 signalling, we generated human monocyte-derived macrophages and challenged them *in vitro* with recombinant Galectin-9 protein (**Figure 6a**). Bulk RNA sequencing revealed a robust transcriptional shift toward an inflammatory phenotype, with upregulation of key cytokines and immune regulators including CCL5, ISG15, CD80, and IL32 (**Figure 6b-d**).

**Figure 6.**
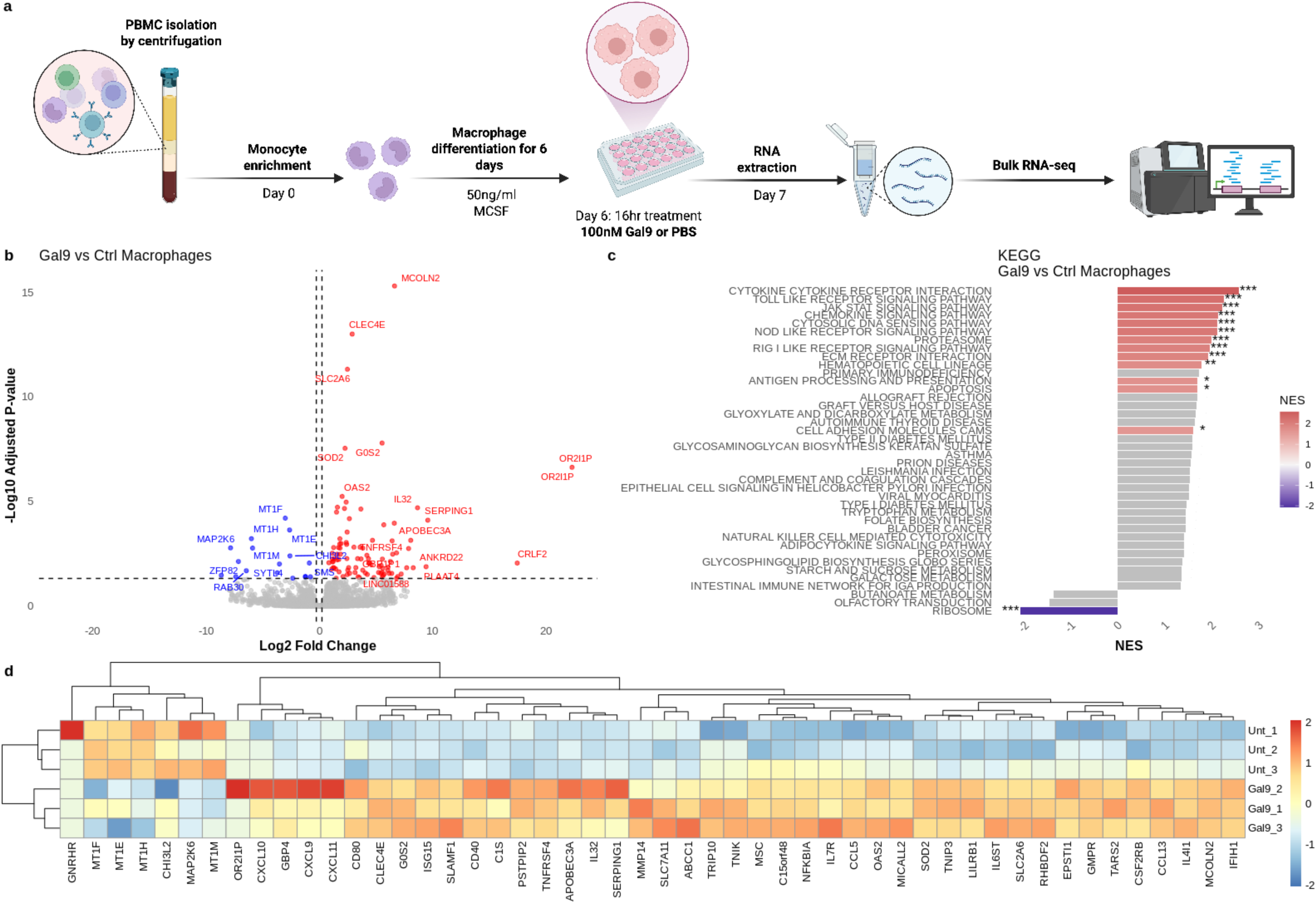
Gene expression changes in human macrophages following Galectin-9 treatment. A. Graphical abstract for isolation, processing, and sequencing of macrophages subjected to Galectin-9 stimulation. B. Volcano plot of Galectin-9 stimulated macrophages vs. control macrophages. C. KEGG Pathway enrichment of Galectin-9 stimulated macrophages, computed by FGSEA using ranked gene statistics; significance stars denote adjusted (p)-value thresholds. D. Top 50 differentially expressed genes (absolute value) between Galectin-9 stimulated and control macrophages.

Collectively, these findings demonstrate that Galectin-1 and Galectin-9 define distinct inflammatory monocyte states with divergent functional programmes in IBD. Galectin-9 further acts as an active proinflammatory stimulus, driving macrophage transcriptional reprogramming toward heightened pro-inflammatory immune activation.

### Serum Galectin-9 levels are elevated in IBD and correlate with inflammatory burden and treatment response

To evaluate whether galectins could serve as a biomarker of intestinal inflammation, we quantified serum Galectin-1, -3 and -9 levels across individuals with CD, UC, healthy controls and symptomatic controls (**Table2 & Figure 7a**). Galectin-9 concentrations were significantly elevated in both CD and UC compared with controls, indicating its association with active intestinal inflammation (**Figure 7a**). Correlation analysis revealed strong positive associations between Galectin-9 and key inflammatory markers, including TNFα and IL6 (**Figure 7b**), as well as broader indices of immune activation such as CRP, total white cell count, and platelet count (**Figure 7c**).

**Figure 7.**
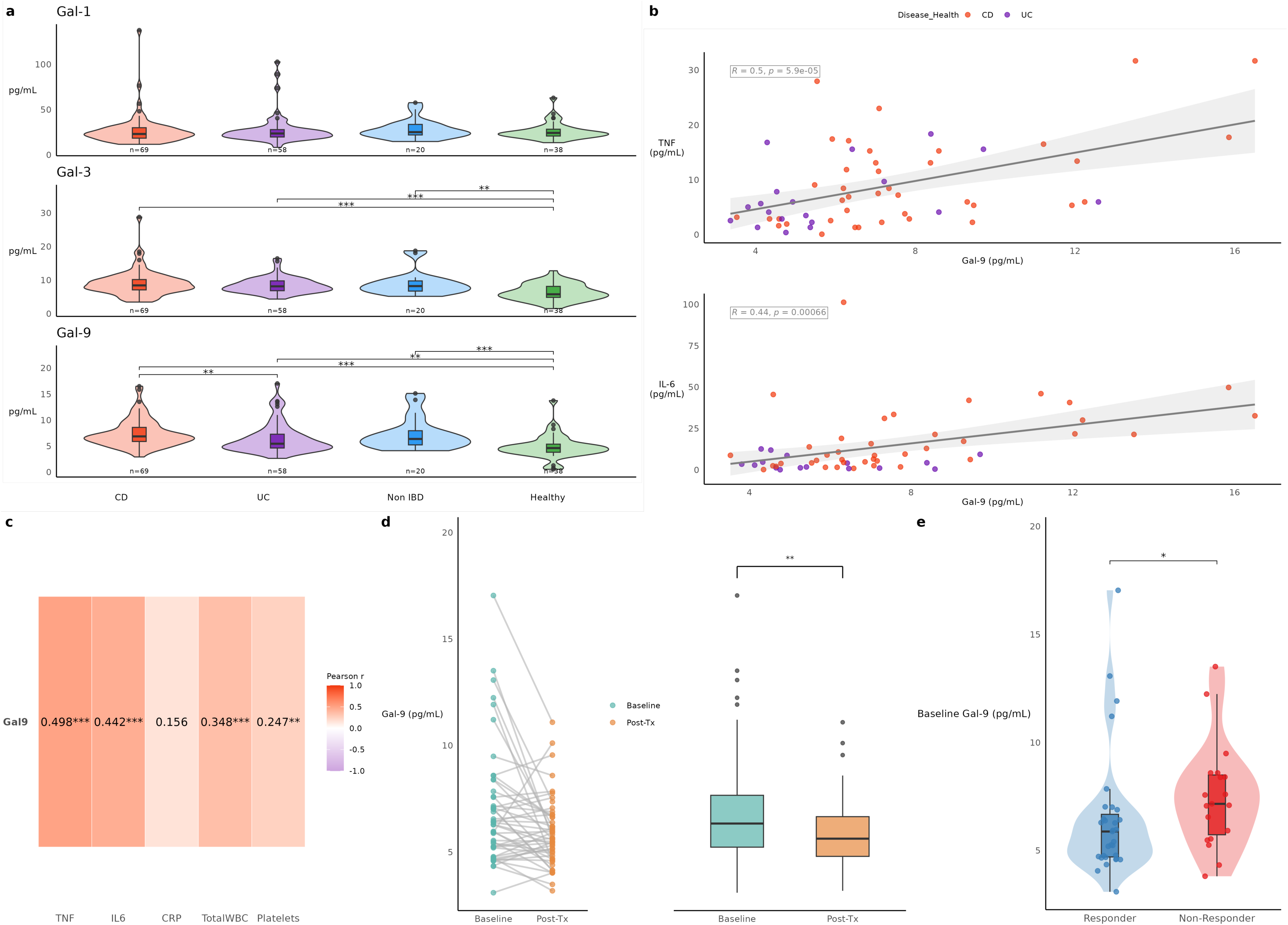
Serum Galectin levels as a marker of disease activity in IBD. A. Serum concentrations of Galectin-1, Galectin-3, and Galectin-9 in patients with ulcerative colitis (UC), Crohn’s disease (CD), non-IBD gastrointestinal disorders (Non-IBD), and healthy controls. Group differences were assessed by pairwise two-sided Wilcoxon rank-sum tests with BH correction. B. Correlation analyses of serum Galectin-9 concentrations with serum TNF-á and IL-6 concentrations in patients with IBD. Associations were assessed using Pearson correlation (with linear fits), shown for CD and UC. C. Pearson correlation coefficients between serum Galectin-9 concentrations and circulating inflammatory markers, including TNF-á, IL-6, C-reactive protein (CRP), total white blood cell count, and platelet count; UC and CD pooled together. D. Serum Galectin-9 concentrations before and after treatment in patients with IBD (CD, n=30; UC, n=26). Patients received standard-of-care therapies, including mesalazine, corticosteroids, immunomodulators, and advanced therapies. Paired baseline vs post-treatment Gal-9 levels were tested using a two-sided paired Wilcoxon signed-rank test. E. Baseline serum Galectin-9 concentrations stratified according to clinical treatment response at 6 months. Baseline responder vs non-responder Gal-9 levels were compared using a two-sided Wilcoxon rank-sum test.

Longitudinal sampling demonstrated that Galectin-9 levels were highest at baseline (visit 1) and declined following treatment (visit 2), mirroring clinical improvement and reduction in inflammatory markers (**Figure 7d**). Patients with higher baseline Galectin-9 levels demonstrated impaired treatment responses at 6 months and were more likely to require advanced therapies during the first year of treatment (**Table 3 & Figure 7e**). These findings position Galectin-9 as a dynamic biomarker of mucosal and systemic inflammation in IBD, with potential utility for disease monitoring and therapeutic stratification.

**Table 3:** Criteria utilised to determine treatment response.

| **Response Grading** | **UC** | **Crohn's** |
| --- | --- | --- |
| **'Partial response' criteria** |  |  |
| Clinical response | $\geq 30\%$ in Partial Mayo<br>**OR**<br>Fall by $\geq 3$ points | Fall in HBI of $\geq 3$ |
| Biochemical response | Fall in FCP $>50\%$<br>**OR**<br>Fall in CRP $>50\%$ | Fall in FCP $>50\%$<br>**OR**<br>Fall in CRP $>50\%$ |
| Mucosal response | Fall in UCEIS $>50\%$ from baseline<br>* If only endoscopic Mayo given, fall $\geq 1$ from baseline | Fall in SESCD $>50\%$ from Baseline<br>**OR**<br>Improvement on imaging from baseline |
| **'Remission' criteria** |  |  |
| Clinical remission | Partial Mayo $<3$<br>No individual score $>1$<br>No steroids or treatment escalation planned or undertaken within 3 months | HBI $<5$<br>No steroids or treatment escalation planned or undertaken within 3 months |
| Biochemical remission | CRP $<5$ AND FCP $<150$ | CRP $<5$ AND FCP $<150$ |
| Mucosal remission | Mayo 0 or UCEIS $\leq 1$ | SESCD $<3$<br>**OR**<br>No active disease on imaging |

HBI = Harvey Bradshaw Index, FCP= Faecal Calprotectin, CRP= C-Reactive Protein, SESCD = Simple Endoscopic Score for Crohn's Disease

## Discussion

This large inception cohort single-cell atlas of newly-diagnosed, adult treatment naïve IBD patients provides a mechanistic basis for understanding the earliest cellular and molecular events that shape disease onset and progression. By profiling one million cells across 137 individuals, this study overcomes a major limitation of previous single-cell analyses, most of which (13, 17, 19, 43, 44), have largely relied on samples from patients with established, treatment-exposed disease. Uzzan *et al.,* (44) also include a cohort of treatment naive patients with UC and previous studies focusing on UC inception cohorts (12, 15, 16) have included small cohorts of treatment-naive patients but there has not been a similar study in adult treatment naive CD patients. Habermann *et al.,* did demonstrate changes in lymphocyte activation and mitochondrial dysfunction in rectal biopsies from treatment-naive children from PROTECT cohort (45, 46). The dataset presented here captures a more homogenous cellular and molecular profile of the patient cohort at presentation by including only treatment-naïve patients and improved single-cell capture [10X source for v3 chemistry improvements], as evidenced by the Bray-Curtis diversity scores. Biologic therapies, corticosteroids, immunomodulators, and 5-ASA compounds are all known to remodel mucosal immunity, alter epithelial-stromal interactions, and reshape the microbiome, thereby obscuring the initiating drivers of inflammation and masking early pathogenic programmes(18).

Smillie et al., previously reported a study of patients largely with established IBD (mostly UC) demonstrating extensive epithelial and stromal remodelling (linked with inflammation associated fibroblasts) and immune activation (inflammatory monocytes, activated T cell etc) (12). They also report crosstalk between immune cells and fibroblasts as a central amplifier of chronic inflammation. Although neutrophils are appreciated as having a central role in IBD, interrogating them using single cell sequencing approaches has historically been technically very challenging. This is due to the very low levels of mRNA present in these cell types and can vary with technologies used with recent data indicating droplet-based approaches like the 10X Chromium used in our study being less amenable (47, 48). Hence, we observed relatively few viable neutrophils from our samples and did not include them in our analysis.

In CD, recent multiomics studies have begun to clarify why a subset of patients develop entrenched, treatment refractory inflammation. Martin *et al*., demonstrated that anti TNFα resistance in CD is driven by a coordinated network of IL1β and OSM producing mononuclear phagocytes, activated OSMR expressing stromal cells, and neutrophil rich inflammatory pathways that together sustain a TNFα independent inflammatory axis in established and active disease (10). Building on this, Kong *et al*., and Thomas *et al*.,(18) profiled both ileal and colonic lesions and mapped longitudinal responses to therapy, showing that anti TNFα treatment remodels immune-stromal interactions only in a subset of patients, while persistent myeloid-fibroblast modules predict nonresponse (16). Collectively, these studies highlight how regional tissue context and progressive remodelling of the mucosal tissue microenvironment shape therapeutic outcomes across IBD. The current dataset therefore offers an unperturbed view of the mucosal microenvironment at the point of diagnosis, enabling the identification of primary inflammatory mechanisms rather than secondary or treatment-induced signatures.

Across both CD and UC, we observed clear and consistent expansion of inflammatory monocytes, along with activated T cells, endothelial cells and IgG-switched plasma cells. These finding align with previous reports of monocyte-driven pathology in established IBD. For example, Huang *et al.,* in a study of patients with UC, CD and unclassified IBD describe an over-expression of pro-inflammatory TNFα expressing macrophages (19). Similarly, Uzzan *et al.,* describe expansion of macrophages with gene signature suggestive of exposure to IgG immune complexes in inflamed compared to uninflamed UC and over-expression in a cohort of UC biopsies from treatment-naive UC patients who turned out to be TNFα inhibitor refractory (44). Our inception cohort demonstrates that these populations are already prominent at disease onset, indicating that they are not just a result of chronic inflammation but early contributors to pathogenesis.

The reduction in compartmental diversity, particularly in the immune compartment, further strengthens the view that inflammation drives towards the expansion of pro-inflammatory subsets. The strong concordance between our dataset and published reference cohorts validates the robustness of our study, while the reduced variability in our inception cohort highlights the importance of studying treatment naïve disease.

To extract disease-relevant gene modules, ligand-receptor cell-cell communication coupled with unbiased tensor decomposition identified the galectin family as being enriched in disease, underscoring the biological relevance of this pathway.

A major output from this current study is the identification of a galectin-centred signalling axis as a dominant, unbiasedly derived signalling module in early IBD. Through CNA and tensor decomp of ligand-receptor matrices, we identified Galectin-1 and Galectin-9 signalling as key drivers of inflammation-associated communications modules. These modules were enriched in inflamed CD and UC and characterised dominant sender-receiver interactions linking inflammatory monocytes, activated CD4⁺ T cells, endothelial cells, and IgG plasma cells. This network architecture suggests that galectin signalling acts as a coordinating hub for mucosal inflammation. Moreover, Galectin-9 and Galectin-1 expressing monocytes were transcriptionally enriched in multiple inflammatory pathways including antigen presentation, immune regulation, migration and microbial sensing. Although the galectins have been implicated in immune regulation and epithelial function (49, 50), they have not previously emerged as central nodes from previous studies in IBD.

As glycan binding proteins, galectins modulate the interface between the colonic mucosa and the microenvironment (microbiota and immune cells), acting to scan and decipher complex cell surface glycan structures and regulating the transition between health (immune tolerance) and disease (chronic inflammation) (51–53).

The potentially pathogenic role of glycans in IBD is well recognised (54) with evidence that glycans structure at inception could act as a prognostic biomarker in UC (55). There has been a small proof of concept trial of altering cell surface glycosylation using topical N acetyl glucosamine to treat refractory proctitis in children (56) with a positive signal of efficacy. Recent data from the GEM study has highlighted serum antiglycan antibodies as a predictive risk factor for developing CD in otherwise healthy first-degree relatives of CD patients (57). Despite the large amount of data concerning the importance of glycobiology in the pathogenesis of IBD and the potential of exploiting this for diagnosis, prognosis and treatment purposes, interventional trials in this space are limited. This is perhaps because many glycans bind multiple receptors, complicating the development of selective drugs in this space (58).

Galectin-1 (LGALS1) functions as a potent immunoregulatory lectin that induces apoptosis of activated Th1 and Th17 cells, stabilises regulatory T cell programmes, suppresses IL-1β and TNF production by myeloid cells, and promotes epithelial restitution through modulation of integrin and cytoskeletal signalling (59, 60). In contrast, Galectin-9 (LGALS9) engages TIM-3 to drive T cell exhaustion preferentially in Th1 and Th17 subsets, enhances monocyte and dendritic cell activation, amplifies IL-1β and TNF dominated inflammatory circuits, and increases endothelial activation and leukocyte (neutrophils/monocytes) adhesion, thereby reinforcing chronic mucosal inflammation(23, 61–63). Analyses focussed on Galectin-9 from heterogenous post-treatment patients with IBD have yielded conflicting data. (64–66). However, a secondary analysis of the pre-treatment RISK cohort highlighted enrichment of Galectin-9 transcripts in inflamed mucosa and PBMCs, positively correlating with disease severity (67, 68).

The translational relevance of these findings is underscored by the serum analyses, which show that circulating Galectin-9 levels are elevated in both CD and UC, correlate with systemic inflammatory markers, and decline following treatment. Galectins are produced by a range of immune and stromal cells. In the context of the healthy gut, Galectin-9 is expressed in epithelial cells (69), resident macrophages (23), dendritic cells (70) and endothelial cells (71) at low levels. During inflammation, expression is significantly increased in activated monocyte-derived macrophages, inflammatory DC and cytokine stimulated T cells, with epithelial and endothelial cells also increasing Galectin-9 production in response to IFNγ, hypoxia and microbial stress (62, 72, 73). Collectively, these cells contribute to the elevated systemic Galectin-9 levels observed, reflecting both local mucosal production and spill-over into the circulation during active inflammation. Previous biomarker studies in IBD have focused on CRP, calprotectin, and cytokines, but few have explored galectins as dynamic indicators of disease activity. Notably, we and others have previously demonstrated that Galectin-9 is also elevated in non-intestinal inflammatory conditions such as peripheral vascular disease, where it tracks with endothelial activation and systemic immune dysregulation, suggesting that Galectin-9 may act as a broader marker of chronic inflammatory burden (74).

Finally, we confirm a direct functional role for Galectin-9 on human monocyte-derived macrophages whereby stimulation of macrophages with Galectin-9 promoted a clear pro-inflammatory signature by bulk RNA-sequencing. These findings reveal that Galectin-9 is not merely a passive biomarker but an active modulator of monocyte and T cell function, reinforcing the mechanistic link between circulating galectin levels and the cellular drivers of early intestinal inflammation. Our data suggest that Galectin-9 reflects both mucosal and systemic inflammatory burden and may have utility for disease monitoring or therapeutic stratification. Given the limited availability of reliable serum biomarkers in IBD, Galectin-9 warrants further evaluation in prospective and interventional cohorts.

An intriguing observation in this study was that the symptomatic control patients had evidence of activated mucosal T cells. Irritable bowel syndrome (IBS) has long been a ‘thorn in the side’ for practising gastroenterologists. The condition is diagnosed from a combination of symptoms corresponding to agreed criteria along with (where available) absence of evident inflammation at endoscopy (75). Previous work has demonstrated the presence of activated T cells in symptomatic control biopsies (12). A recent systematic review and meta-analysis of IBS studies have reported that IBS can be distinguished from asymptomatic healthy individuals based on elevated circulating cytokines and faecal calprotectin (76). The symptomatic non-inflamed controls in our study all had at least a transient elevation of faecal calprotectin. Further longitudinal study of such individuals may well lead to important clinical insights.

In summary, the findings of this study position galectin centred signalling as a unifying inflammatory axis in early IBD, driven predominantly by transcriptionally distinct monocyte subsets. This work reshapes our understanding of the initiating events in IBD pathogenesis and highlights galectin mediated pathways as potential therapeutic targets or prognostic biomarkers. Future studies should investigate whether modulation of galectin signalling can attenuate mucosal inflammation, whether galectin defined monocyte states predict treatment response, and how these pathways interact with microbial and stromal cues during disease evolution. By establishing a high resolution, treatment naïve single cell atlas of IBD onset, this study provides a foundational resource for mechanistic discovery and therapeutic innovation in IBD.

### Grant Support

The research was carried out at the National Institute for Health and Care Research (NIHR) Birmingham Biomedical Research Centre (BRC). The work is supported by multiple funding including, F. Hoffmann-La Roche Ltd. AJI Supported BHF Project grant PG/23/11476 and Birmingham Fellowship. AAM extends his appreciation to the Deanship of Research and Graduate Studies at King Khalid University for funding via small group research project under grant number RGP1/82/46.

### Disclosures

These authors disclose the following: Matthew Leipner, Alexandra Paun, Virginie Sandrin, Patrick Trenkle, Achim Klein, Sabrina Danillin and Daniel Regan-Komito are employed by F. Hoffmann-La Roche, Ltd. The remaining authors disclose no conflicts.

### Data Transparency Statement

The processed, anonymized, and cell-type-annotated single-cell and gene expression matrices supporting the findings of this study have been deposited in Zenodo under DOI: 10.5281/zenodo.21026153 (Version 1.0.0). The dataset is currently held under an embargo period to protect proprietary discovery phases during peer review. Reviewers and editors may access the private files securely using the provided journal access token. The underlying raw sequencing data are not publicly available due to commercial confidentiality and patient privacy. Detailed computational analysis pipelines, workflow instructions, and specific software versions are fully documented in the Supplementary Methods section.

## Author Contributions

- **Conceptualization:** VS, DRK, AJI
- **Methodology:** VS, TI, DRK, AJI
- **Investigation:** ML, PR, ST, AP, JB, AAM, AS, NS, JC, FM, PT, AK, SD
- **Resources (clinical samples):** ML, PR, ST, AP, JB, AAM, AS, NS, JC, FM, PT, AK, SD, TI
- **Data Curation:** ML, PR, ST, AP, JB, AAM, AS, NS, JC, FM, PT, AK, SD
- **Formal Analysis:** ML, PR, ST, AP, JB, AAM, AS, NS, JC, FM, PT, AK, SD
- **Writing – Original Draft:** ML, PR, TI, AJI, DRK
- **Writing – Review & Editing:** All authors
- **Supervision:** TI, DRK, AJI

## Abbreviations

CCC: Cell-Cell Communication
CD: Crohn’s Disease
CNA: Co-Varying Neighbourhood Analysis
DEG: Differentially Expressed Gene
FDR: False Discovery Rate
GSEA: Gene Set Enrichment Analysis
IBD: Inflammatory Bowel Disease
IBS: Irritable Bowel Syndrome
MAD: Median Absolute Deviation
PBMC: Peripheral Blood Mononuclear Cell
SC: Single Cell
UC: Ulcerative Colitis
UMI: Unique Molecular Identifier

