## Supplementary Methods for "Single-cell analysis of an adult IBD INCEPTION cohort reveals Galectin-linked disease mechanisms"

### 2.1 Patient Enrolment and Biopsy Collection

All diagnoses were established using history, biochemistry, endoscopy, histological and radiological criteria in line with the European Crohn's and Colitis Organisation guidelines [1]. Clinical, endoscopic and patient reported outcome measures were collected at baseline and longitudinally during the first year of treatment. Subsequent treatment was not protocolised, but therapeutic outcomes, including the need for surgery, were collected prospectively. Those where IBD and other significant organic diseases (colorectal cancer, immune mediated disease, other colitis e.g. ischaemic) were excluded during this process formed a symptomatic control (SC) cohort.
For serum samples, 12ml of blood was collected in clot activator and separation gel tubes. After 15-30 mins, samples were centrifuged at ~1000g for 15 minutes. Separated serum was mixed by inversion and divided into 1ml aliquots. Samples were frozen at -80°C within 1.5 hours of collection until analysis.

### 2.4 $CD14^+$ $CD16^-$ classical monocyte isolation

Classical ($CD14^+CD16^-$) monocytes were isolated from PBMCs by the initial depletion of $CD16^+$ cells. PBMCs were incubated with CD16 human microbeads (Miltenyi) (50ml per 50 $x10^6$ cells) for 20 minutes at 4°C and subsequently passed through an LD magnetic column (Miltenyi) to eliminate $CD16^+$ cells. The negative fraction was then labelled with CD14 human microbeads (Miltenyi) (50ml per 50 $x10^6$ cells) for 20 minutes at 4°C and processed through an LS magnetic column (Miltenyi). The positively selected fraction containing $CD14^+CD16^-$ cells were collected and counted as previously mentioned.

### 2.7 Single Cell RNA Sequencing

**Single-cell RNA sequencing data processing**

Single-cell RNA sequencing data were processed using a Snakemake v8.30.0 workflow implementing single-cell best practices as compiled by the Theis laboratory [2, 3]. Raw sequencing data were aligned and quantified using 10x Genomics Cell Ranger v9.0.1 against the GRCh38-2024-A human reference transcriptome (GENCODE release 40) [4].

**Ambient RNA removal, quality control, and cell filtering**

To correct for ambient RNA contamination, CellBender v0.3.2 was applied to raw count matrices from Cell Ranger output [5]. Further filtering and quality control was performed on an individual sample basis using a multi-step filtering strategy implemented in Scanpy v1.11.4 [6, 7]. Initial filtering removed low-quality cells with fewer than 100 UMI counts or fewer than 150 detected genes. Each sample was then clustered via leiden clustering, and median absolute deviation (MAD)-based outlier detection was then applied for multiple quality metrics on a per-cluster basis to account for biological heterogeneity:

- Total UMI counts (outliers by log-transformed counts, max cutoff: 4,000)

- Number of detected genes (min cutoff: 200 genes)
- Percentage of counts in top 20 genes
- Mitochondrial gene content (max cutoff: 50%, with cells exceeding 80% flagged as extremely high)

Doublet detection was performed using scDblFinder v1.20.2 implemented in R v4.4.3 on filtered count matrices to compute doublet scores and classify cells as singlets or doublets [8].

**Cell type annotation**

Automated cell type annotation was performed using CellTypist v1.7.1 with multiple reference models [9]:

1. "Cells_Intestinal_Tract" model (based on Elmentaite et al.) [10]
2. "Adult_Human_Intestine" model (trained on Smillie, Elmentaite, Burclaff, and James datasets) [10]
3. Pan-GI Atlas "Adult/Paediatric Full Healthy Reference" model [11]
4. Pan-GI Atlas "Adult/Paediatric Large Intestine" model [11]

Cell type labels were hierarchically organized into three levels (celltypist_labels1-3) and mapped to tissue compartments (epithelial, immune, stromal) using a custom mapping file. CellTypist majority voting was used to refine initial predictions, and confidence scores were retained for quality assessment.

**Data integration and batch correction**

Individual samples were merged into one dataset and subsequently processed by tissue compartment. For each compartment, highly variable genes (HVGs) were identified using the Seurat v3 method (n = 3,000 genes, span = 1) with batch-aware selection across samples of Scanpy. Integration and batch correction were performed using scVI v1.0.1 (single-cell Variational Inference) from scvi-tools, a deep generative model that learns a low-dimensional representation of the data while accounting for technical batch effects [12]. The scVI model was configured with:

- 2 latent layers
- Covariate encoding (batch, mitochondrial percentage, ribosomal percentage)
- Deep covariate injection disabled
- Layer normalization enabled for both encoder and decoder
- Batch normalization disabled

Models were trained on GPU (CUDA v12.4.1) with PyTorch v2.0.0.post200 and automatic stopping criteria [13, 14] . Dimensionality reduction was performed on the scVI latent space using UMAP (Uniform Manifold Approximation and Projection), and clustering was performed using the Leiden algorithm (resolution = 1.0) with GPU-accelerated neighborhood graph construction via RAPIDS (k = 8 neighbors) [15, 16, 17].

**Single Cell RNA Sequencing Co-Varying Neighborhood Analysis**

Co-varying neighbourhood analysis (CNA) was used to identify cell populations and microenvironmental contexts associated with disease state using the cna Python package [18]. CNA identifies local cell neighborhoods whose abundance varies systematically with sample-level phenotypes while controlling for technical and biological confounders.

Data from batch-corrected, compartment-specific subsets were converted to MultiAnnData format, with sample-level metadata organized in a dedicated dataframe. Cell-cell similarity graphs were computed on the scVI latent representations using GPU-accelerated neighbor finding (RAPIDS, k neighbors). For each analysis, samples were restricted to a single disease cohort (e.g., Crohn's disease patients only) to focus on within-disease heterogeneity.

Sample-level phenotypes were encoded as binary variables for CNA analysis:

- Disease state: inflamed vs. non-inflamed (based on endoscopic assessment)
- Sex: male vs. female
- Batch: encoded from Arvados collection identifiers

CNA association testing was performed using the `cna.tl.association()` function with the following specifications:

Phenotype of interest: inflammation status (inflamed vs. non-inflamed)

Covariates: sex (to control for sex-related differences)

Batch effects: Arvados collection batch (to account for technical variation in sequencing batch)

The CNA algorithm constructs a Neighborhood Abundance Matrix (NAM) by computing, for each cell, the representation of its local neighborhood in terms of sample membership. Principal component analysis is performed on the NAM to identify major axes of cross-sample variation in neighborhood composition. For each neighborhood (cell), CNA computes a neighborhood correlation score (ncorr) quantifying the association between the local cellular environment and the phenotype of interest, while regressing out effects of covariates and batch.

Statistical significance was assessed using permutation testing with false discovery rate (FDR) correction. Neighbourhoods with significant associations (FDR < 0.05) were retained for downstream analysis. Neighborhood correlation scores were categorized as:

- **Expanded** (ncorr > 0.2): neighborhoods enriched in inflamed samples
- **Depleted** (ncorr < -0.2): neighborhoods enriched in non-inflamed samples
- **Neutral** (|ncorr| ≤ 0.2): neighborhoods without strong association

Neighborhood correlation scores computed by CNA were stored with single-cell metadata and used to stratify cells in downstream analyses. Specifically, cells were categorized by their ncorr values to examine whether disease-associated versus control-associated neighborhoods exhibit different Cell-cell communication patterns (integrated with LIANA+ analysis). This approach enabled identification of disease-relevant cell types and their interactions within inflamed microenvironments.

**Cell-cell communication analysis**

Cell-cell communication analysis was performed using LIANA+ v1.0.4 (Ligand-Receptor Analysis) on compartment-specific datasets [19]. Results were aggregated across multiple methods using LIANA's rank aggregation approach, combining magnitude and specificity rankings across most frequently used cell-cell communication tools inlcuding [CellPhoneDB, CellChat, SingleCellSignalR] using the CellPhoneDB, CellTalkDB, CellChatDB, and OmniPath databases [20, 21, 22, 23, 24].

**Tensor decomposition**

To identify coordinated patterns of cell-cell communication across samples, tensor decomposition was performed using cell2cell v0.7.4 and tensorly v0.8.1 [25, 26]. LIANA+ communication scores (magnitude_rank) were aggregated into a 4-dimensional tensor with dimensions representing: (1) samples (n contexts), (2) ligand-receptor pairs, (3) sender cell types, and (4) receiver cell types. Communication scores were transformed using an inverse function (1 - x) to convert ranks to weights, with outer joining across all samples.

The optimal number of factors (latent components) was determined through elbow analysis evaluating reconstruction error across ranks 1-15. Non-negative tensor factorization was performed using the PARAFAC (Canonical Polyadic) decomposition method implemented in tensorly with PyTorch backend for GPU acceleration [27]. The factorization was configured with:

- Random initialization (random_state = 42)
- Regular optimization mode (100 maximum iterations, tolerance = $10^{-7}$)
- GPU computation (CUDA v12.4.1)
- Rank = 5 factors based on elbow analysis

Factor activities across disease conditions were compared using Mann-Whitney U tests with Benjamini-Hochberg false discovery rate correction ($\alpha = 0.05$). Samples were grouped by disease status (e.g., CD-Inflamed, CD-Non-Inflamed, UC-Inflamed, UC-Non-Inflamed, Non-IBD) for statistical comparisons.

**Pathway Enrichment Analysis**

To interpret the biological meaning of tensor factors, pathway enrichment analysis was performed on ligand-receptor factor loadings. Gene Set Enrichment Analysis (GSEA) was conducted using ligand-receptor gene sets derived from KEGG pathway annotations [28]. LR pairs were mapped to pathways based on their constituent genes, with a minimum set size of 15 genes. GSEA was performed with 999 permutations (random_state = 6) and significance threshold of adjusted p-value ≤ 0.05. Normalized Enrichment Scores (NES) were computed to identify pathways enriched (NES > 0) or depleted (NES < 0) in each factor.

Additionally, pathway activity footprints were estimated using PROGENy (Pathway RespOnsive GENes) gene sets through multivariate linear models (MLM) [29]. PROGENy gene sets (top 5,000 genes, human organism) were intersected with ligand-receptor pairs to estimate pathway activities for each factor. Activities were visualized using hierarchical clustering with z-score normalization across factors.

Cell-cell communication networks were visualized for each factor by computing joint loadings as the product of sender and receiver cell factor loadings. Interactions with joint loadings above a threshold of 0.075 were visualized as directed networks, with node sizes proportional to cell type importance and edge weights representing communication strength.

### 2.9 Serum Enzyme-linked Immunosorbent Assays (ELISA)

ELISA were undertaken on serum obtained from CD, UC and SC at baseline and longitudinally. For each ELISA, stored serum was thawed on ice and vortexed during this process and again just prior to aliquoting. Serum was pre-aliquoted into 96-well plates for dilution as per the manufacturer's instructions for a given ELISA. Standard protocols for each ELISA were followed and assays performed for Gal-1, Gal-3, Gal-9, TNF, IL-6 and IL-23 (Quantikine ELISA, R&D systems, Biotechne). Standard curves and samples were run in duplicate wherever possible. Optical density was determined by reading the plates at different wavelengths. Readings were taken at 450 and 540nm using a Biotek Synergy HT microplate reader and Kc4 analysis software within 30 minutes of stopping the ELISA. The coefficient of variation (CV) between each duplicate was calculated for the 450nm reading and again for the delta value to ensure data was consistent. The delta values were then averaged, and the average of the blank value from the standard curve subtracted.

### 2.10 Statistical Analysis

For ELISA data, concentration for the samples was interpolated from the standard curve using a four-parameter logistic curve fit in Prism (GraphPad, California, US). Values above the upper limit of the standard curve were obtained using a simple linear regression. Results were corrected for initial dilution to generate the data used in the analyses. Non-parametric testing was used for onward analysis. For two independent data sets, a Mann Whitney U was utilised with a standard p-value threshold of 0.05. A Wilcoxon-signed-rank was used for two paired groups. For tests including more than two independent groups, a one-way ANOVA (Kruskal-Wallis test by ranks) was used, with a non-parametric repeated measures ANOVA (Friedman test) for paired groups. P values for such analyses are presented after correction using the Holm method to control for the family wise error rate.

**Computational environment**

All analyses were performed using Python v3.10.12 with Scanpy v1.9.3, AnnData v0.9.1, NumPy v1.24.4, pandas v1.5.3, scVI v1.0.1, decoupleR v1.5.0, and LIANA v1.0.4 [30, 31, 32]. Visualization was performed using matplotlib v3.7.1 and seaborn v0.11.2. Celltypist and scDblFinder were run using R v4.4.3. GPU-accelerated computations utilized RAPIDS with CUDA v12.4.1 and PyTorch. The complete workflow was processed using Snakemake on a high-performance computing cluster with both CPU and GPU resources. Analysis notebooks were executed using Jupyter Lab v4.1.6 [33, 34, 35, 36, 37].

**Supplementary Methods References**

29. Van Rossum G and Drake FL. Python 3 Reference Manual. CreateSpace. 2009.
30. R Core Team. R: A language and environment for statistical computing. R Foundation for Statistical Computing. 2021. https://www.R-project.org/
31. Badia-i-Mompel P, Vélez Santiago J, Braunger J, et al. decoupleR: ensemble of computational methods to infer biological activities from omics data. Bioinformatics Advances. 2022;2:vbac016. https://doi.org/10.1093/bioadv/vbac016
32. Harris CR, Millman KJ, van der Walt SJ, et al. Array programming with NumPy. Nature. 2020;585:357-362. https://doi.org/10.1038/s41586-020-2649-2
33. McKinney W. Data structures for statistical computing in Python. Proceedings of the 9th Python in Science Conference. 2010:56-61.
34. Pedregosa F, Varoquaux G, Gramfort A, et al. Scikit-learn: Machine learning in Python. Journal of Machine Learning Research. 2011;12:2825-2830.
35. Waskom ML. seaborn: statistical data visualization. Journal of Open Source Software. 2021;6:3021. https://doi.org/10.21105/joss.03021
36. Hunter JD. Matplotlib: A 2D graphics environment. Computing in Science & Engineering. 2007;9:90-95. https://doi.org/10.1109/MCSE.2007.55
37. Kluyver T, Ragan-Kelley B, Pérez F, et al. Jupyter Notebooks – a publishing format for reproducible computational workflows. Positioning and Power in Academic Publishing: Players, Agents and Agendas. 2016:87-90.
